# Accounting for pseudo-replication of Linkage Disequilibrium for contemporary *N_e_* estimation

**DOI:** 10.64898/2026.09.09.750346

**Authors:** Tin-Yu J. Hui, Laiyin Zhou, Austin Burt

## Abstract

The Linkage Disequilibrium (LD) of unlinked loci can be used to estimate contemporary effective population size (*N_e_*) of one to a few generations ago. In genomic datasets loci on different chromosomes are considered unlinked, but there are many more pairs of unlinked loci than there are independent pairs of chromosomes, resulting to confidence intervals (C.I.) being too narrow if the non-independence is not taken into account. Simulations were run to investigate the correlation structure among LD of unlinked loci, which can be expressed by the LD of loci along the same chromosomes, based on a discovery of a novel Random Probe LD estimator. We classify the correlation into two categories: overlapping of loci and disjoint pairs. The former is induced from the same locus being considered twice and is the stronger form of correlation. These correlations feed into *ρ*, a parameter to quantify the degree of pseudo-replication in a dataset, and further a correction formula from which C.I. can be properly inferred. We demonstrate the use of our method via an analysis of genomic data from the malaria-transmitting *Anopheles gambiae s.s* mosquitoes. Apart from the point and C.I. estimates, we find that 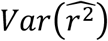 is inflated by about 550 times due to pseudo-replication, highlighting the danger of not handling genetic correlation properly.

## Introduction

The effective population size (*N*_*e*_) is a key genetic parameter that quantifies the magnitude of drift and the loss of heterozygosity. Depending on the questions of interest, some studies focus on the longer-term *N*_*e*_, spanning hundreds of thousands of generations, or across multiple sub-populations to cover a wider geographical area. On the other end of the spectrum, such as for the purposes of conservation and population monitoring, the relevant measure should be the recent *N*_*e*_, which also tends to be a local measure as fewer migration events are captured (Wang and Whitlock, 2003). The consensus is that a minimum *N*_*e*_ of 500 is required to maintain the balance between mutation and the loss of additive variations, bur a larger threshold is usually preferred to prevent mutational meltdown. An instantaneous bottleneck can drastically reduce genetic diversity, which requires a very long time for it to recover. Beyond stochastic evolution, *N*_*e*_ also determines the relative importance of migration and selection (Wang, 2005).

There are two main ways to estimate recent *N*_*e*_ from genetic markers: the temporal methods, which track the changes in allele frequency over time (Nei and Tajima, 1981; Williamson and Slatkin, 1999; Hui and Burt, 2015; Waples et al., 2025), and the linkage disequilibrium (LD) methods, estimated between unlinked loci (Hill, 1981; Waples and Do, 2010). The former requires sampling from two or more time points to estimate the harmonic mean *N*_*e*_ over the sampling horizon. In contrast, the one-sample LD method estimates the *N*_*e*_ of the parental generation, although a pair of unlinked loci may have a decaying memory about *N*_*e*_ of up to three to four generations (Waples, 2005).

The theories behind LD, drift, and recombination have been studied extensively (Hill and Robertson, 1968; Sved and Feldman, 1973; McVean, 2002; Park, 2012). In short, the expected standardised correlation of gene frequency *E*(*r*) approaches 0 due to non-zero recombination, but its variance *Var*(*r*) = *E*(*r*^2^) > 0 is a function of the recombination rate *c* and *N*_*e*_ (Hill, 1981). Unlinked pairs (*c* = 0.5, i.e. those with 50% chance of recombining per generation) have short memory on population histories thus are best to estimate the most recent *N*_*e*_, while linked loci reflect past population sizes as a weighted average with exponential-like memory decay (Hayes et al., 2003; Santiago et al., 2020). Estimation of *r*^2^ relies on the identification of haplotype frequencies. For unphased (genotypic) data, likelihood-based or gene counting methods are required to resolve the double heterozygotes (Weir, 1979; Excoffier and Slatkin, 1995; Waples, 2006). Besides, most observed *r*^2^ is biased upward due to finite diploid sample size *s* (*s* << *N*_*e*_ for most cases). It is generally thought that the observed *r*^2^ can be partitioned into two components: that induced by sampling and evolutionary forces (Waples, 2006; Hui and Burt, 2020).

Pseudo-replication arises when observations or summary statistics are not truly independent (Waples et al., 2021). In genomics, for instance, neighbouring loci tend to evolve together and thus carry correlated information. This results in the amount of information the data conveyed being less than nominal, that we might fall into the trap of being too confident when constructing confidence intervals (C.I.). Pseudo-replication was not a big issue decades ago when data contained merely a handful of genetic loci or microsatellites, but it can no longer be ignored in the genomic era when hundreds of thousands of tightly linked loci can be sequenced or simulated. Such an effect is particularly pronounced in the estimation of LD as it is based on pairs of loci. This study aims to investigate the correlation structure among unlinked *r*^2^ through computer simulations, then proposes a correction for proper C.I. estimation. We largely follow the mathematical notations used by previous studies for continuity (Table 1).

**Table 1:** Mathematical notations used in this study.

| Symbol | Description |
| --- | --- |
| $N_e, \widehat{N_e}$ | Effective population size and its estimate |
| $r^2, r_{ij}^2$ | LD between a pair of alleles from two loci, often comes with subscripts to indicate the loci of interest |
| $\widehat{r^2}$ | The overall LD of the observed dataset, includes both drift and sampling components |
| $s$ | Diploid sample size |
| $c$ | Recombination rate between a pair of loci, $c = 0.5$ for unlinked loci |
| $L_1, L_2$ | Number of loci on chromosomes 1 and 2 |
| $\rho$ | A measure of pseudo-replication |
| $C$ | Number of chromosomes, $C \geq 2$ |

### Theories and Methods

The classical scenario, as assumed by LDNe and NeEstimator (Do et al., 2014), considers a system of *L* mutually unlinked loci (Figure 1, top panel), which holds for organisms with numerous autosomes or genetic loci that are distant from each other. This is becoming very restrictive for organisms with very few pairs of autosomes (such as Anopheles gambiae, see also our worked example below), or the more general scenario of a genomic dataset for genomic datasets with densely linked loci along the same chromosome, or in To illustrate the concept, this work focuses on a two-chromosome diploid population with biallelic SNPs (Figure 1B). SNPs on the same chromosome are linked with various degrees of linkage, only pairs from opposite chromosomes are guaranteed to be unlinked. We can use all *L*_1_*L*_2_ unlinked pairs to get the point LD estimate:

**Figure 1.**
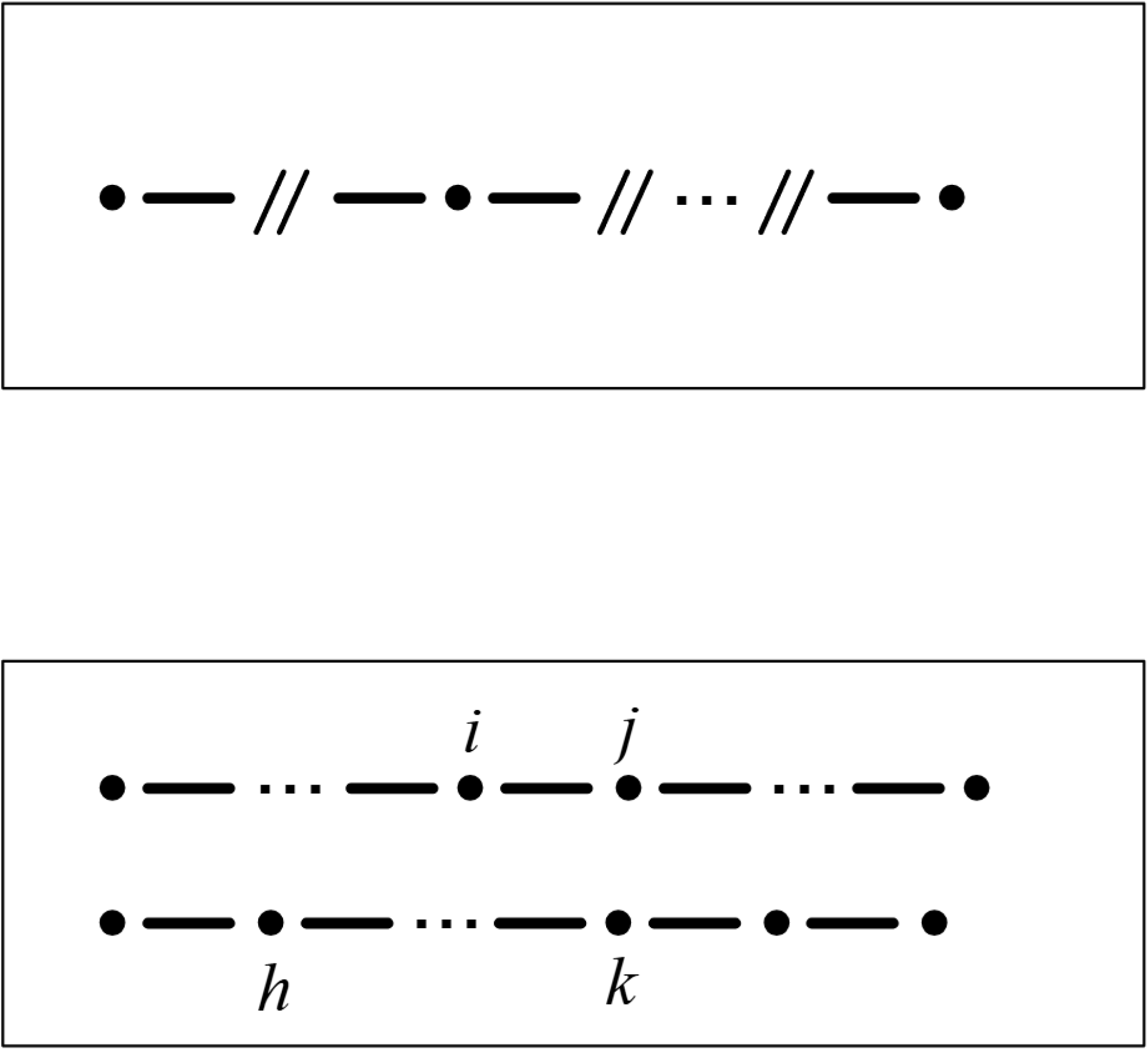
Schematic diagrams for the two scenarios of unlinked LD. [Above] The mutually unlinked case, as discussed by Do et al. (2014) and most previous studies. Between every loci (dots) is a recombination breakpoint (double-slash) with rate *c* = 0.5. [Below] The focus of this work concerning a typical genomic dataset with two (or more) chromosomes. Loci on the same chromosomes (e.g. *i* and *j*) are linked with various degrees of recombination. Pairs from opposite chromosomes (e.g. *j* and *h*) are guaranteed unlinked.

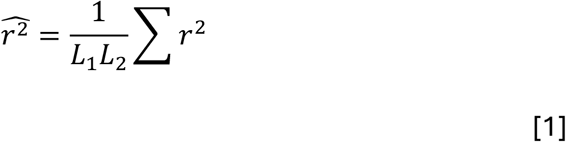

where *L*_1_ and *L*_2_ are the number of SNPs on the two chromosomes respectively. With unphased genotypes, unlinked 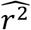 is usually estimated via the Burrows’s method (Weir, 1979?) with the Waples’ empirical formulae (Waples, 2006) for converting the overall 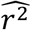 into 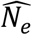. Note that Equation 1 still holds even when the *r*^2^ are correlated, but the sample variance depends on the pairwise correlation:

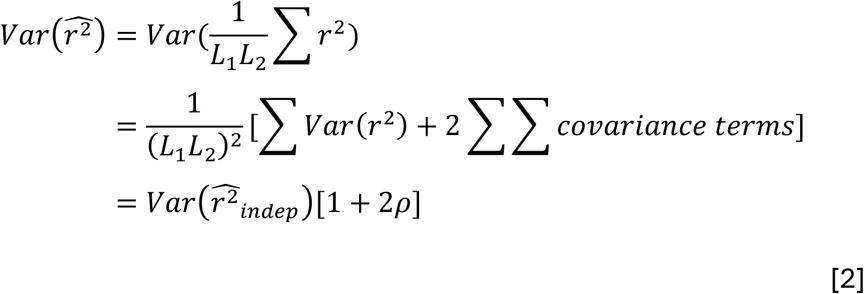

where

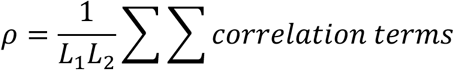

Equation 2 on the second moment is exact. Note that there are *L*_1_*L*_2_(*L*_1_*L*_2_ − 1)/2 correlations terms in *ρ*, which quantifies the severity of pseudo-replication by measuring how much the variance is inflated compared to 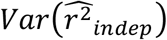, which is the sampling variance if all *r*^2^ are assumed to be independent.

To better understand *ρ* and the correlation terms let us consider the following system of four loci: *i, j* on the first chromosome, and *h, k* on the second (Figure 2), yielding four unlinked *r*^2^ for point estimation via Equation 1. They come with six correlation terms, all contributing to *ρ*. The stronger form of correlation comes from “overlapping of pairs”, referring to the same locus being involved in both *r*^2^, for example, locus *i* in 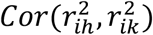. The second form comes from “disjoint pairs” involving four different loci, such as 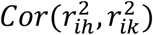, which is presumably weaker (Figure 2).

**Figure 2.**
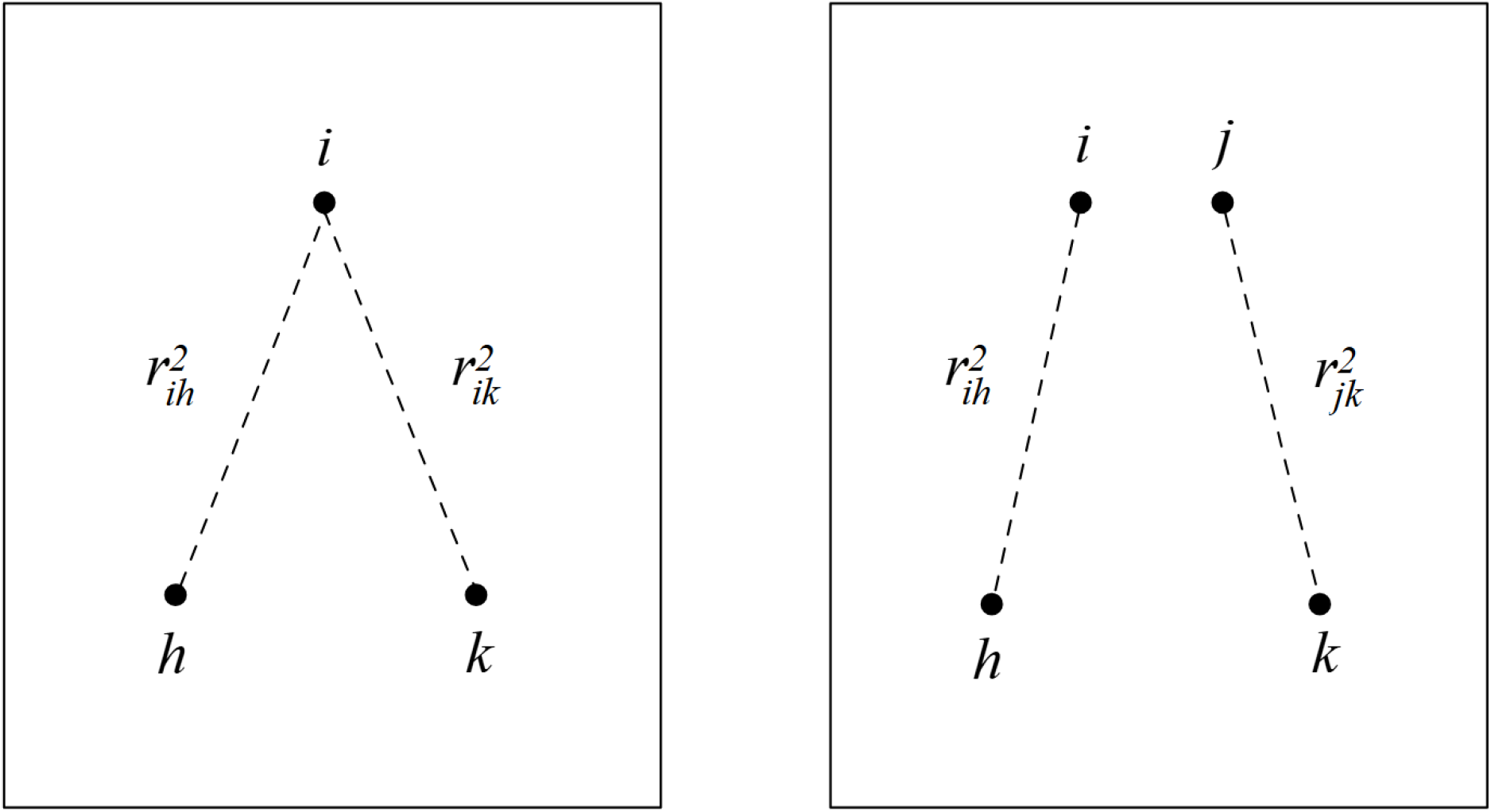
The two forms of correlation concerning the unlinked *r*^2^ from a genomic dataset. Loci *i* and *j* are on one chromosome, and *h* and *k* on another (see Figure 1, bottom panel). [Left] Overlapping of loci. Locus *i* contributes to both the *r*^2^ between (*i* and *h*) and (*i* and *k*). As a result, 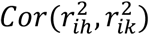 is found to be quite high. [Right] Disjoint pairs. The two unlinked *r*^2^ involves four distinct loci.

What exactly are these correlation terms? In a separate study, Hui and Burt (in prep) proposed the novel Random Probe (RP) LD estimator, that the *r*^2^ between a pair of loci can be estimated from the correlation of *r*^2^ between each of the two loci with a set of randomly generated loci. We learn that the RP estimator is largely unbiased, thus

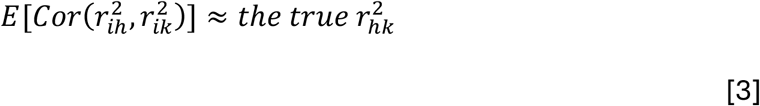

where locus *i* on the opposite chromosome is considered as one of the random probe loci. Here the “true” component refers to the population *r*^2^ which is unaffected by sampling (i.e. solely induced by evolutionary forces, specifically drift in this study). This accounts for the correlation from the overlapping of pairs.

### Simulations

While extensive simulations had been run in the development and verification of the RP estimator, here we built an individual-based Wright-Fisher forward simulator for our four-locus system specifically for the context of pseudo-replication. The average *r*^2^ along the same chromosome from 5,000 independent simulations are shown in Figure 3. On each chromosome, the two loci were in complete initial LD (i.e. 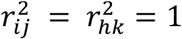) before drift and recombination acted on the pair to cause LD decay over time. We also calculated 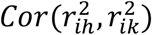 and 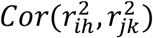 at each time point from repeated simulations and plotted them on the same graph. Figure 3 verified our claim in Equation 3 on the correlation measure, and further

**Figure 3.**
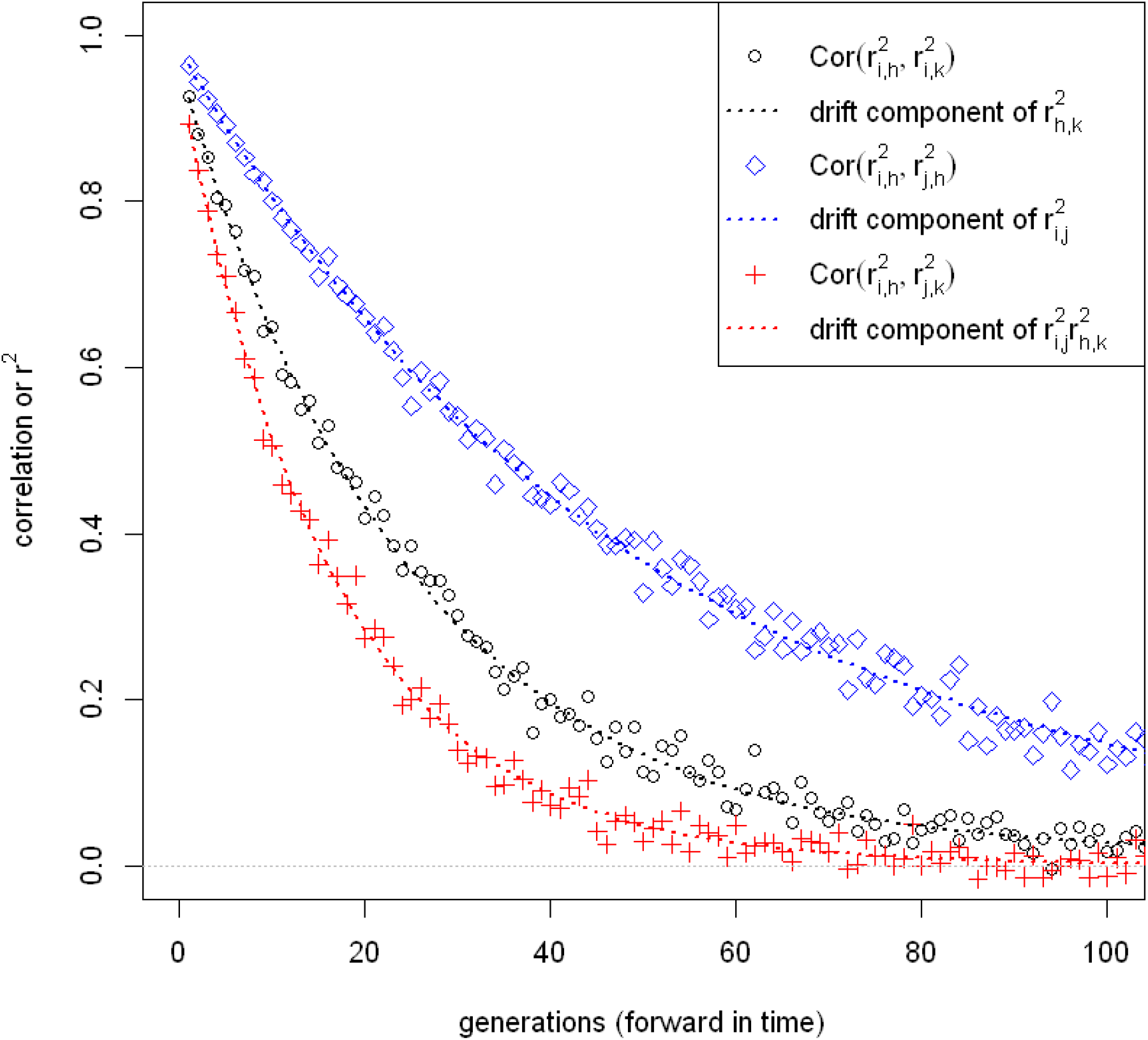
Simulation results for the 4-locus system (Figure 2). The aim here is to evaluate the correlation between pairs of unlinked LD. With 5,000 independent sets of simulations, the sample Pearson correlations 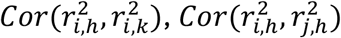, and 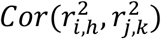 were calculated at each time point, and are shown as diamonds, dots and crosses respectively. The first two correlations are examples of overlapping of loci, while the last one (crosses) arises from disjoint pairs. From the same set of simulations, we plotted the average drift (true) components of 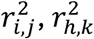, and 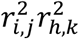 as dotted lines. Loci on the same chromosome are in complete LD initially, with recombination rate of *c* = 0.01 between loci *i* and *j*, and *c* = 0.02 between *h* and *k* on the second chromosome. True *N*_*e*_ = 1000 with diploid sample size *s* = 200. See Figure S1 for full simulation results.

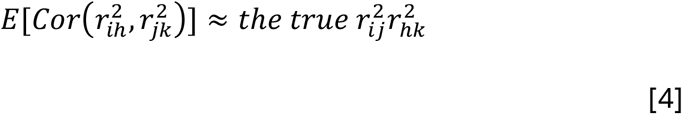

In other words, the correlation among unlinked *r*^2^ can be expressed as the LD (or product of two LD) induced by drift along the same chromosome. Since everyone knows how to estimate LD from a pair of loci, these correlations can now be resolved. Note that Equation 3 is a special case of Equation 4 as 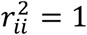. Similar results were also found in other simulation settings (S.I.).

With the correlation terms and *ρ* clarified, the remaining question is to find 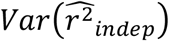, the variance of the point estimate when all unlinked *r*^2^ are (wrongly) assumed to be independent. Classical result suggests that 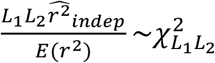 for *L*_1_ *L*_2_ unlinked pairs, which has a variance of 2*L*_1_*L*_2_ (Waples, 2006). Therefore, in our simulations with 4 unlinked pairs, the ratio of the observed 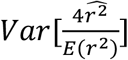 to that of 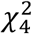 should closely follow (1 + 2*ρ*), where *ρ* is estimated from the pairwise correlation terms (Equations 3-4, Figure 4).

**Figure 4.**
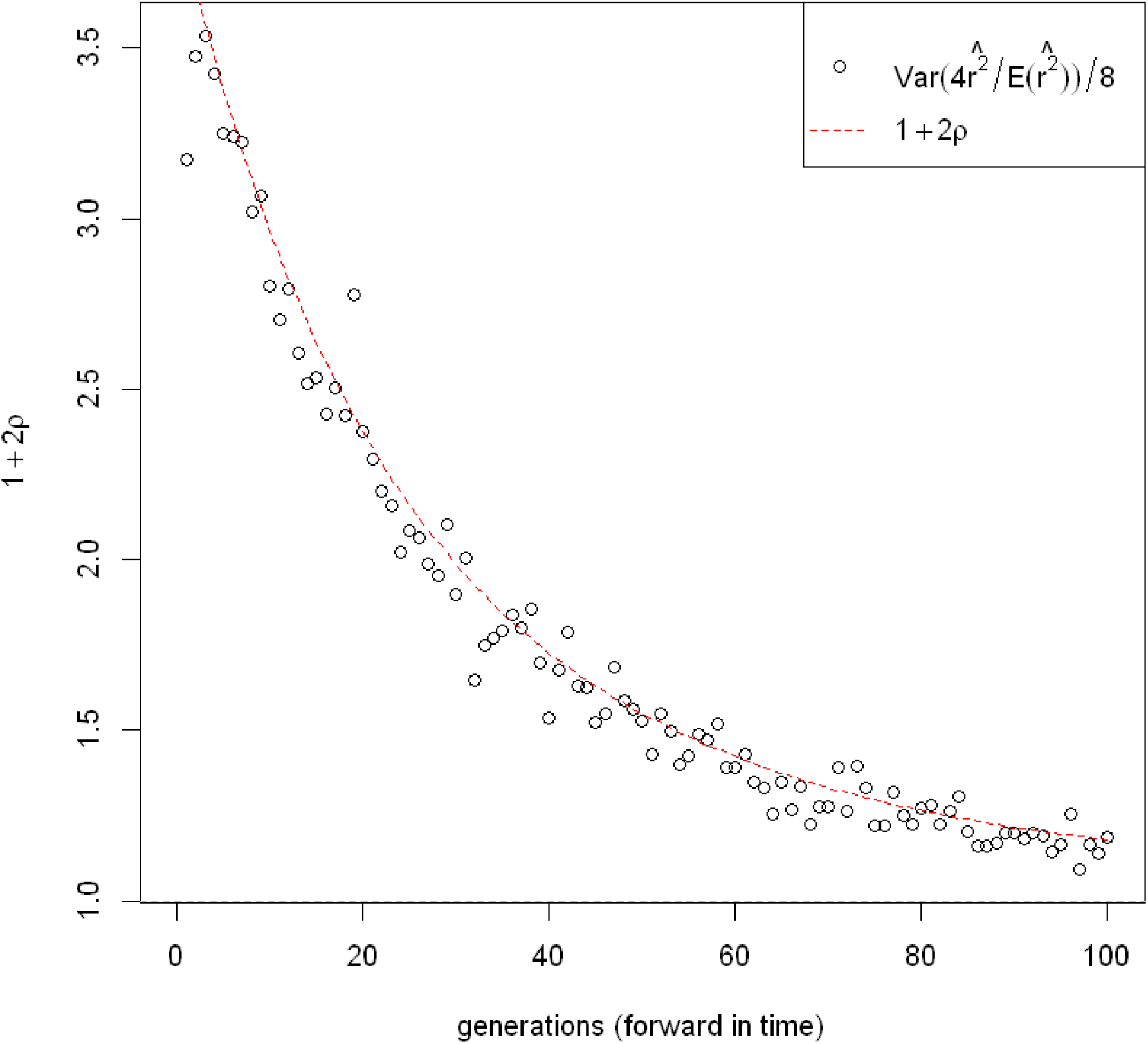
Plot of (1 + 2*ρ*) over time in the same system as in Figure 3. The average (1 + 2*ρ*) calculated from the pairwise correlation terms via Equation 2 from 5,000 independent simulations (red dashed line), alongside the empirical 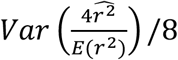 from the same simulations (dots). The denominator 8 comes from the variance of 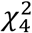.

The C.I. is constructed based on the same argument, that with pseudo-replication the empirical 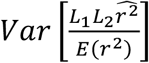 follows of 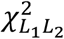, multiplied by a factor (1 + 2*ρ*):

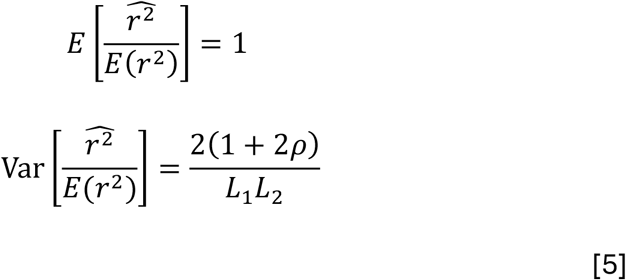

Normal approximation holds for large *L*_1_ *L*_2_, and thus the (1 − *α*) C.I. for 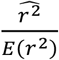 is 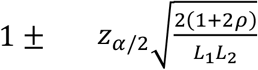, where *z*_*α*/2_ is the *α*/2 quantile of the standard normal. Putting everything together:

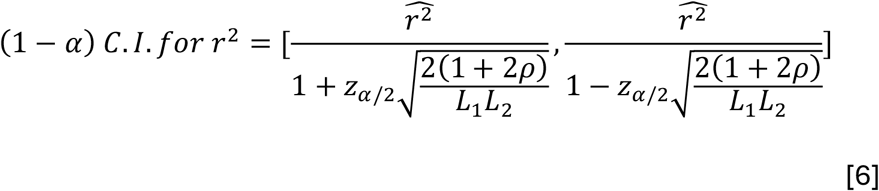

### Applications

We analysed a whole-genome sequencing dataset from the MalariaGEN Vector Observatory, an international collaboration working to build capacity for malaria vector genomic research and surveillance. The dataset consists of 68 *An. gambiae (s*.*s*.) mosquitoes collected from Kiimi, an island on Lake Victoria in Uganda, between May and June 2014. Four-fold degenerate SNPs, which are considered genetically neutral, were extracted from the two autosomes. SNPs with missing alleles or minor allele frequency (maf) of <10% were removed. A two-step LD pruning was performed to avoid tightly linked loci along the same chromosome: 1 SNP was chosen per 500bp non-overlapping window, then followed by sequential LD pruning to remove pairs with *r*^2^ > 0.35. After filtering, 5,368 and 3,552 biallelic SNPs from chromosome arms 2R and 3R were retained and entered the calculation. The point estimate from all the unlinked pairs from opposite chromosomes was 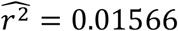, equated into 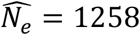 via Waples’ formulae. For C.I., we followed the procedure to compute *ρ* ≈ 275, and the 95% C.I. for *N*_*e*_ = [662, 9707] after correction. The empirical distribution of 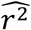 can be visualised in Figure 5, alongside the uncorrected one for comparison.

**Figure 5.**
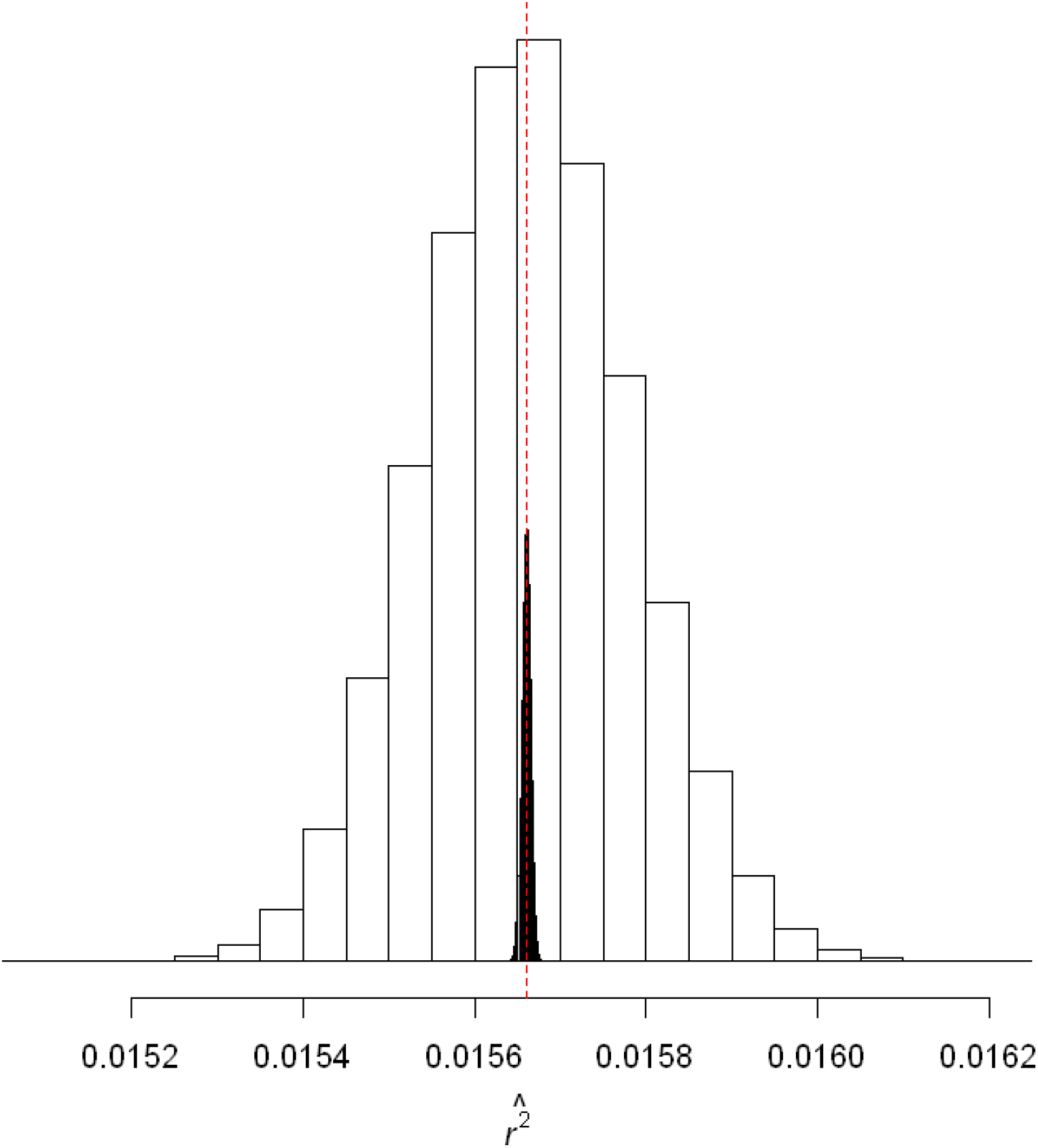
The sampling distribution of 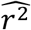 from our worked example, shown as the white histogram based on normal approximation. It was based on the genotypes from 68 *An. gambiae* collected from Kiimi island, Uganda. Please refer to the main text for sampling and SNP ascertainment. The unlinked *r*^2^ and point *N*_*e*_ estimate were computed via the Waples’ formulae with the Burrows’ method, while the *r*^2^ along the same chromosome were estimated via the method by Ragsdale and Gravel (2019). The point estimate 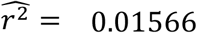 is highlighted by the red dotted line, which corresponds to 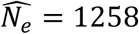. The unadjusted distribution (i.e. assuming all unlinked *r*^2^ are independent) is shown in grey, which is much narrower.

## Discussion

Jackknife, as its name suggests, is a drop-in method to mitigate pseudo-replication in genomics and beyond (Bhatia et al., 2013). While no knowledge on the underlying distribution is required, it is unclear on which dimension (loci? Individuals? or the *L*_1_*L*_2_ unlinked *r*^2^?) jackknife should be applied in the context of LD estimation (Jonas et al., 2016). The pseudo-values from the minus-one jackknife are almost certainly non-independent thus it cannot provide a proper variance estimate. Waples et al. (2016) discussed using datasets of similar type to estimate recent *N*_*e*_ but from a different approach: The authors mixed linked loci with unlinked loci to infer recent *N*_*e*_, which were known to be biased downward because *E*[*r*^2^] ≈ 1/(1 + 4*N*_*e*_*c*) (Sved and Feldman, 1973). Note that the *r*^2^’s from various *c* respond to past population sizes differently as weighted averages. The same paper proposed an empirical bias correction, however, only under the additional assumption of constant *N*_*e*_, which would have been very different if there were fluctuating *N*_*e*_ or seasonal dynamics. In this work we use only truly unlinked pairs from different chromosomes for point estimation in Equation 1 to ensure past population histories do not interfere with the contemporary *N*_*e*_ estimate.

While only *r*^2^ from unlinked pairs contribute to the point estimate, no information goes to waste as those from linked loci along the same chromosomes quantify the magnitude of pseudo-replication through *ρ*. When dissecting the correlation among unlinked *r*^2^, one immediate challenge is dimensionality, that the number of *r*^2^ is a quadratic function of *L*, and further the number correlation terms in Equation 2 is quartic! We classified the correlation into one of the two forms: overlapping of loci and disjoint pairs of *r*^2^. Higher correlation is anticipated when the same locus appears in both *r*^2^, especially when the other two on the same chromosome are already in high LD. Despite deviating from its original application, the expected correlation due to overlapping of loci (Equation 3) is a direct consequence of the RP estimator (Hui and Burt, in prep) by treating the locus on the opposite chromosome as a probe locus. Individually speaking, correlation from disjoint pairs is considerably smaller, but the aggregated effect is no less important as there are *L*_1_*L*_2_(*L*_1_ − 1)(*L*_2_ − 1)/2 of them (note: there are *L*_1_*L*_2_(*L*_1_ + *L*_2_ − 2)/2 from overlapping of loci, which is a factor of about *L*/2 smaller, assuming *L*_1_ = *L*_2_ = *L*). Unless the average LD on the same chromosome is smaller 2/*L* (which is unlikely given large *L* in genomic datasets), correlation from disjoint pairs continues to contribute to *ρ*.

Temporal *F*, another widely used estimator for contemporary *N*_*e*_, had also suffered from pseudo-replication until recently. *F* is calculated on per locus basis, thus a correction was implemented to decorrelate drift signals at nearby loci (Hui et al., 2021). The current work is the LD analogue to that for temporal *F* but with a different correlation structure. Note that LD, being a single-sample estimator for *N*_*e*_, does not require knowledge of recombination rates (or recombination maps) along the chromosomes, which adds another advantage over temporal *F*. The effect of pseudo-replication is stronger in smaller populations (Waples et al., 2021) in which tighter C.I. is needed most for decision making. We applied our method to estimate the contemporary *N*_*e*_ for a genomic dataset of *An. gambaie*, the primary malaria vector in Uganda (alongside *An. arabiensis* from the same species complex). Our estimate was comparable to other published figures on the same and nearby islands on Lake Victoria, while the *N*_*e*_ of the mainland populations were previously reported to be much larger (Wiltshire et al., 2018; Mwima et al., 2026). The census size (estimated separately from close-kin mark-recapture, for example) could be at least an order of magnitude higher due to sweepstake-like reproductive success (Lerch et al., submitted), thus studying *N*_*e*_ alone may underestimate epidemiological indicators for malaria exposure and human-mosquito interaction. Another interesting result from our worked example is the *ρ* estimate and the implied degrees of freedom. Although thousands of loci from two chromosome arms were involved, they contained the roughly same information as 34,604 independent pairs of unlinked loci (5,368*3,552/(1+2*275)). While it is difficult to make a direct comparison between the two *N*_*e*_ estimators, LD computes on pairs of loci thus in theory should experience a much higher degree of pseudo-replication. The concept can be generalised to an arbitrary number of chromosomes *C* by considering all 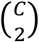 pairs of chromosomes in-turn. All else being equal, the number of unlinked *r*^2^ increases with *C*, and further the proportion of the correlation caused by overlapping of loci decreases. These explain the patterns observed by Waples et al. (2021) that weaker pseudo-replication is associated with more chromosomes.

Our model assumes one closed population of diploid individuals. The population reproduces according to the Wright-Fisher model with random mating and discrete generation (Do et al., 2014). LD arises from drift, while other forces, such as linked selection and migration, are assumed absent. There have been numerous discussions on the practical aspects such as on bias correction, missing alleles, and maf filtering (Waples, 2006; Peel et al., 2013), and these results should large apply to this work. LD pruning is an iterative process to remove loci in high LD along the same chromosome. We see very little value to include loci in high LD, as they do not give much additional signal towards *N*_*e*_, while the extra sampling noise is likely to be detrimental. More stringent criterion to filter for missing alleles and maf can be applied given the vast number of loci. In our worked example, we first selected one locus from each 500bp non-overlapping window, based on a typical LD decay pattern of *An. gambiae* (Ag1000G, 2017), to coarsely remove loci in tight linkage, before conducting another round of LD pruning on the chosen subset. We intentionally avoided inversion-rich regions (e.g. 2La) which are known to suppress local recombination. Most analyses were performed in R (R Core Team, 2025), while some components from the simulation and the worked example were coded in C and CUDA for performance.

## Conclusions

With the advance in sequencing technologies, obtaining large number of linked loci (e.g. from whole-genome sequencing) is increasingly affordable and feasible even for non-model species. *N*_*e*_ continues to play a vital role in conservation and population monitoring as it has recently been incorporated as one of the headline indicators by the Convention on Biological Diversity (Thurfjell et al., 2022). It is essential to ensure the existing *N*_*e*_ estimators (or summary statistics in general), many of which were developed in the pre-genomic era, will continue to perform as expected. Methods to mitigate pseudo-replication can be as empirical as jackknife, nonetheless, additional insight is always welcome by dissecting the correlation structure as in the case we did for temporal *F* (Hui et al., 2021). Together with this work, we have refreshed the two most popular methods for contemporary *N*_*e*_ estimation for genomic datasets.

## Acknowledgements

We thank Rita Mwima for useful insights on the geography of the Ugandan islands. All authors receive funding through Target Malaria, which is supported by grants from the Gates Foundation and Coefficient Giving. This work is supported by Wellcome Trust (224487).

## Data availability

Computer codes are found in https://github.com/tinyuhui/LD_pseudo_replication

---

**Figure S1.**
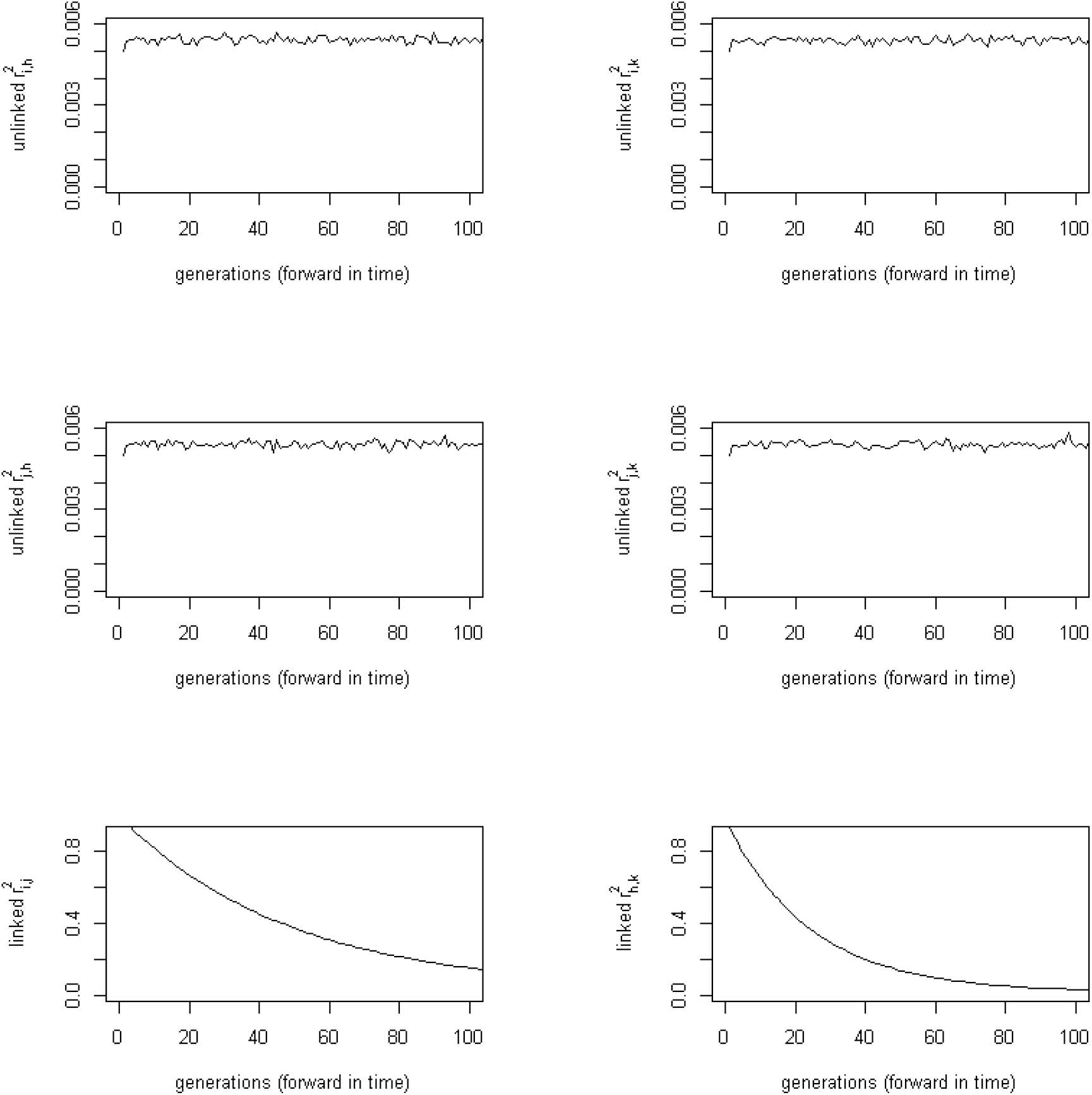
Plots of all six Burrows’ *r*^2^ over time in the same four-locus system (Figures 3-4). There are four unlinked *r*^2^ from opposite chromosomes (top and middle rows) which have the expectation of 0.0054 under *N*_*e*_ = 1000 and *s* = 200 according to the Waples’ empirical formulae. Unlinked pairs have very short memory thus they reach their equilibrium in almost no time. The bottom row displays the two *r*^2^ from linked pairs along the same chromosomes with gradual decay over generations. The recombination rate between loci *h* and *k* is 0.02 (bottom right), twice as large as another pair, thus 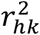 has a faster decay (and a lower equilibrium value).

